# Pediatric Sarcoma Data Forms a Unique Cluster Measured via the Earth Mover’s Distance

**DOI:** 10.1101/116384

**Authors:** Yongxin Chen, Filemon Dela Cruz, Romeil Sandhu, Andrew Kung, Prabhjot Mundi, Joseph Deasy, Allen Tannenbaum

## Abstract

In this note, we combined pediatric sarcoma data from Columbia University with adult sarcoma data collected from TCGA, in order to see if one can automatically discern a unique pediatric cluster in the combined data set. Using a novel clustering pipeline based on optimal transport theory, this turned out to be the case. The overall methodology may find uses for the classification of data from other biological networking problems.

## 1 Introduction

The present note describes a novel method for data clustering applied to the classification of pediatric sarcoma data. Namely, in this work, we combined two data sets: the first consisting of the gene expression of predominantly pediatric sarcoma patients, and the second consisting of the gene expression of adult sarcoma patients taken from the The Cancer Genome Atlas (TCGA) database. We then wanted to see if one could discern some quantifiable difference between the pediatric and adult cases.

Accordingly, we applied a method based on the Earth Mover’s Distance (EMD) to the data; see Sections 2 and 4 below for all the details. Briefly, the proposed pipeline constructs a weighted graph based on the network topology inferred from the Human Protein Reference Database (HPRD), and then treating the graph as a Markov chain, constructs the invariant (stationary) measure, computes the pairwise distances via EMD among all the networks, and then represents the resulting distance matrix as a heat map. Other than an outlier (see Section 3 below), our method was able to segregate the pediatric cases from the adult cases, i.e., we found two rather distinct clusters.

We should note that ideas based on the Earth Mover’s Distance (also known as the Wasserstein 1-metric [19, 26, 27]) have already been applied in studying various properties of cancer networks. In particular, the Wasserstein 1-metric leads to a notion of curvature [17] that turns out to be positively correlated with network robustness; see [4, 20]. This geometric network approach to studying cancer, led to some work indicating that cancer networks are more functionally robust than their normal counterparts [20].

The EMD (and more generally optimal mass transport theory) is very natural for studying the properties of various weighted graphs modeling biological networks, since it gives a natural metric between probability distributions. Its use has become very widespread in recent years being employed for problems in communications, finance, engineering, and biology [6, 19, 26, 27]. This work continues this line of research, by using the distance to cluster biological data.

Finally, we believe that the overall pipeline can be more generally applied in clustering many different types of network data (represented as a weighted graph). We note that we associate the invariant measure to each individual network in the class of data to be classified, and then apply the EMD. This is a distinct advantage since no preprocessing is necessary, other than normalizing the weighted graphs to ensure that they define a Markov process [13]

## 2 Results

### 2.1 Data

The gene expression data sets used in the present work, consist of two parts. The first part includes the gene expression of 27 patients diagnosed with pediatric-associated sarcoma and treated at Columbia University Medical Center (CUMC). Informed and signed consent for clinical and research sequencing was obtained in the context of the pediatric precision medicine program (PIPseq) established at CUMC and under the CUMC Institutional Review Board (IRB)-approved protocol AAAN8404 [15]. The second part was downloaded from The Cancer Genome Atlas (TCGA) database, covering the gene expression data of 265 adult patients. We have one sample per patient for both of them, so 292 samples in total. The data sets were normalized utilizing one of the standard methods for treating RNA-Seq counts data via the variance-stabilizing transformation (VST) in the DESeq2 package for R [14]. This normalization was done amongst all of the 292 sarcoma samples.

The network topology (graph adjacency matrix) was constructed using interaction information from the Human Protein Reference Database (HPRD) [18]. Specifically, we took the intersection of the genes that appear in both HPRD data and the gene expression data, and then kept the largest connected component. After discarding the redundant genes, we arrived at a gene regulatory network with 8844 nodes (genes) and 34926 edges (interactions). The average and median degrees are 7.9 and 4, respectively.

### 2.2 Weighted graph and invariant measure

We constructed a weighted graph for each sample using the mass action principle [25]. In particular, for given gene expression {*x_i_ >* 0 | 1 ≤ *i* ≤ *n*} the weight *p_ij_* on the edge (*i*, *j*) is defined as

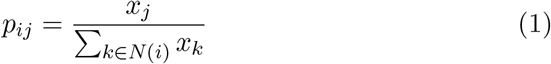

for any *j* ∈ *N*(*i*). Here *n* = 8844 is the number of nodes and *N*(*i*) denotes the set of neighbors of the node i. Note by construction the matrix 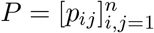 is a stochastic matrix and satisfies that *p_ij_* = 0 if the edge (*i*, *j*) doesn’t exist.

The stochastic matrix *P* defines a Markov chain [13] on the gene regulatory network. Different properties such as entropy and curvature have been considered for this object to study robustness of cancer network [20, 29]. Here we consider the invariant measure (stationary distribution) of this Markov chain. The Markov chain describes the information flow between genes. When the underlying network is connected, the system will eventually reach an equilibrium and this equilibrium is described by the invariant measure. Mathematically, it is a probability vector π satisfying

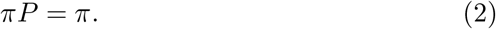

Thus π is a left eigenvector of *P* with non-negative entries that sum to 1. The value π_*i*_ at node *i* reflects the portion of contribution of that node to the entire network. In other words, the invariant measure π is a centrality measure of the significance of different genes.

In general, to obtain the invariant measure, one needs to solve the linear equation (2). However, for the specific stochastic matrix in (1), π has the explicit structure

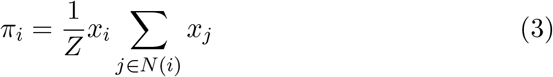

where *Z* is a normalization factor (partition function) forcing π to be a probability vector.

The expression (3) is very interesting. Note that the value of π_*i*_ at node *i* reflects the significance of gene *i* in the gene regulatory network. It consists of two components: the gene expression level *x_i_* of gene *i* and the total gene expression of its neighbors ∑_*j∈N*(*i*)_*x_j_*. ***In other words, the invariant measure captures the key property that a gene is important if its expression level is high and it interacts with many other genes.***

### 2.3 Optimal transport on graphs

Consider a connected undirected graph 𝒢 = (𝒱, ε) with *n* nodes in 𝒱 and *m* edges in ε. Given two densities *ρ*^0^*ρ*^1^ ∈ ℝ^*n*^ on the graph, the original formulation of the optimal transport problem seeks a joint distribution μ ∈ ℝ^*n×n*^ of *ρ*^0^ and *ρ*^1^ minimizing the cost the total cost ∑*c_ij_μ_ij_* that is,

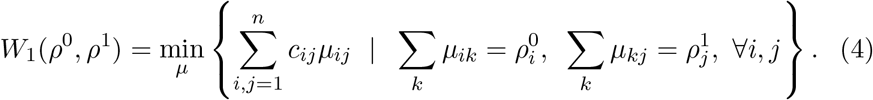

Here *c_ij_* is the cost of moving unit mass from node *i* to node *j* and is taken to be the minimum of the number of steps to go from *i* to *j*. For example, if the edge (*i*, *j*) exists, then *c_ij_* = 1. The minimum of this optimization problem defines a metric *W*_1_ (the Earth Mover’s Distance) on the space probability densities on 𝒢. An alternative formulation is defined on the fluxes *u* ∈ ℝ^*m*^ on the edges. Let *D* ∈ ℝ^*n×m*^ be the oriented incidence matrix of 𝒢, then

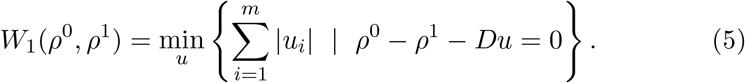

Note that the incidence matrix *D* = [*d_ik_*] ∈ ℝ^*n×m*^ is defined by associating an orientation to each edge *e_k_* = (*i,j*) = (*j*, *i*) of the graph: one of the nodes i, j is defined to be the head and the other the tail, and then we set *d_ik_* = +1(–1) if *i* is the head (tail) of *e_k_* and 0 otherwise. Compared to (4), which has *n*^2^ variables, the above formulation has only *m* variables. It may greatly reduce the computational load when the graph 𝒢 is sparse, i.e., m << *n*^2^. This is the case in our data sets, where *n* = 8844 and *m* = 34926. In implementation, we used the standard convex optimization package CVX [3] written in Matlab, in order to numerically solve (5).

### 2.4 Clustering of sarcoma data

We define a distance function between different gene expression data sets using optimal transport theory on graphs. More specifically, we define the distance between two gene expression data sets to be the *W*_1_ optimal mass transport distance between the two invariant measures induced by the gene expressions as in (3). This distance *W*_1_ can be computed through convex programming [6, 19]. We computed the *W*_1_ distances between each pair of all the 292 samples (27 pediatric sarcoma and 265 adult cancer). The heat map of the resulting distance matrix is as shown in Figure 1. The samples clearly split into two clusters; one cluster for the 27 pediatric sarcoma samples and one cluster for the 265 adult cancer patients.

**Figure 1:**
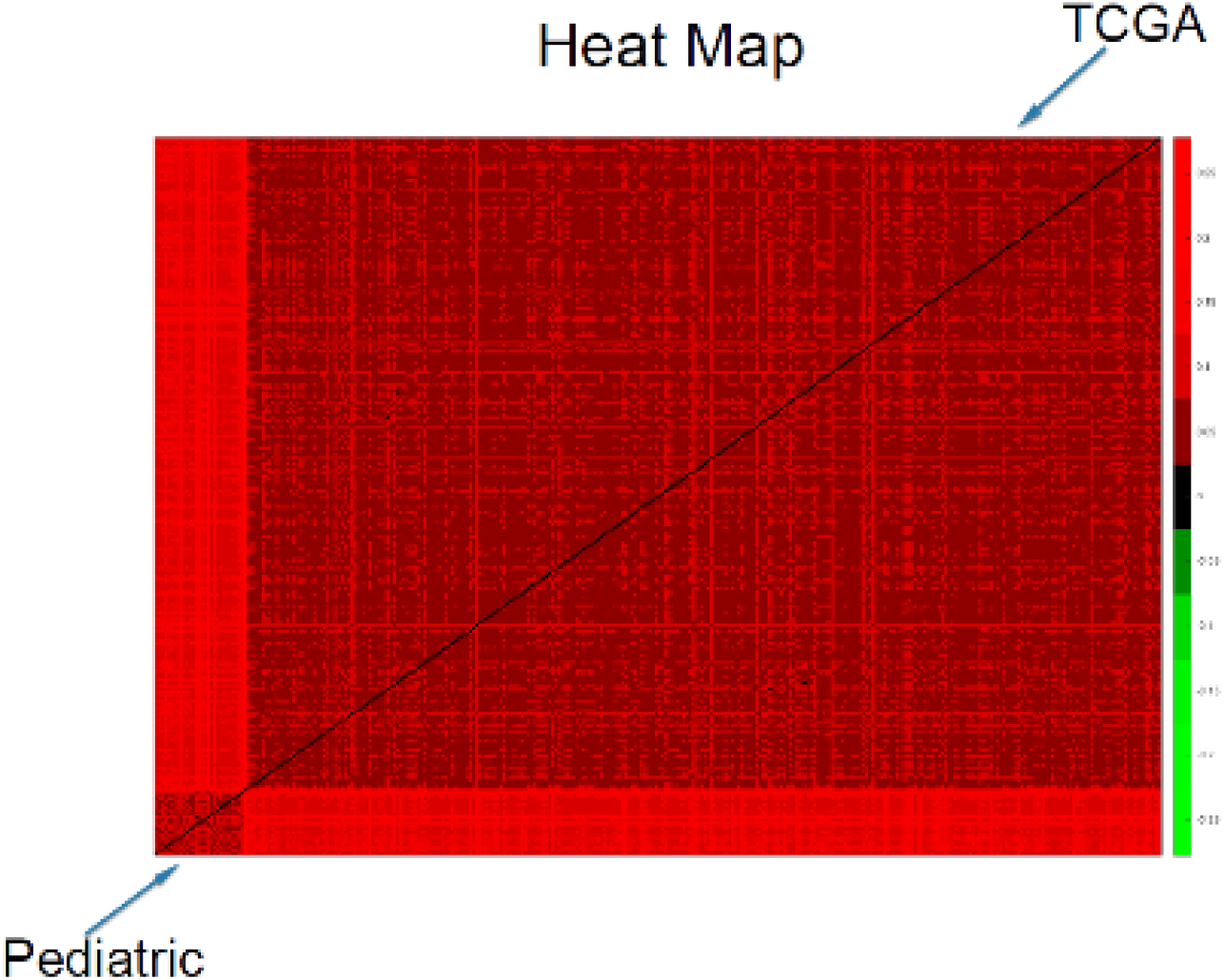
Heat Map Showing Pediatric Cluster

To visualize more clearly the two clusters, we truncate the distances using some threshold: set the value to be zero if the distance is less than the given threshold and one otherwise. The results with threshold value o.o75 and 0.1 are depicted in Figure 2 and 3, respectively. Note that there is a small gap between these two clusters, which indicates that the last sample in the pediatric sarcoma is an outlier. Figure 4 is a 3D plot of the distance matrix, from which we can see an obvious difference that distinguishes this outlier from the rest of the sarcoma samples. The clusters and the outlier can be also seen based on the histograms. Figures 5–7 are the histograms of the distances within the pediatric sarcomas, within the adult sarcomas, and between these two age groups, respectively. Apparently the distances within the two groups (pediatric, adult) are smaller than the distances between them. In particular, the average distances within the two groups are 0.0891, 0.0665 while the average distance between them is 0.1366. The distance between the outlier and the other samples is shown in Figure 8, with mean value 0.2424, which is significantly larger than the average. See our discussion in the next section for further analysis of these results.

**Figure 2:**
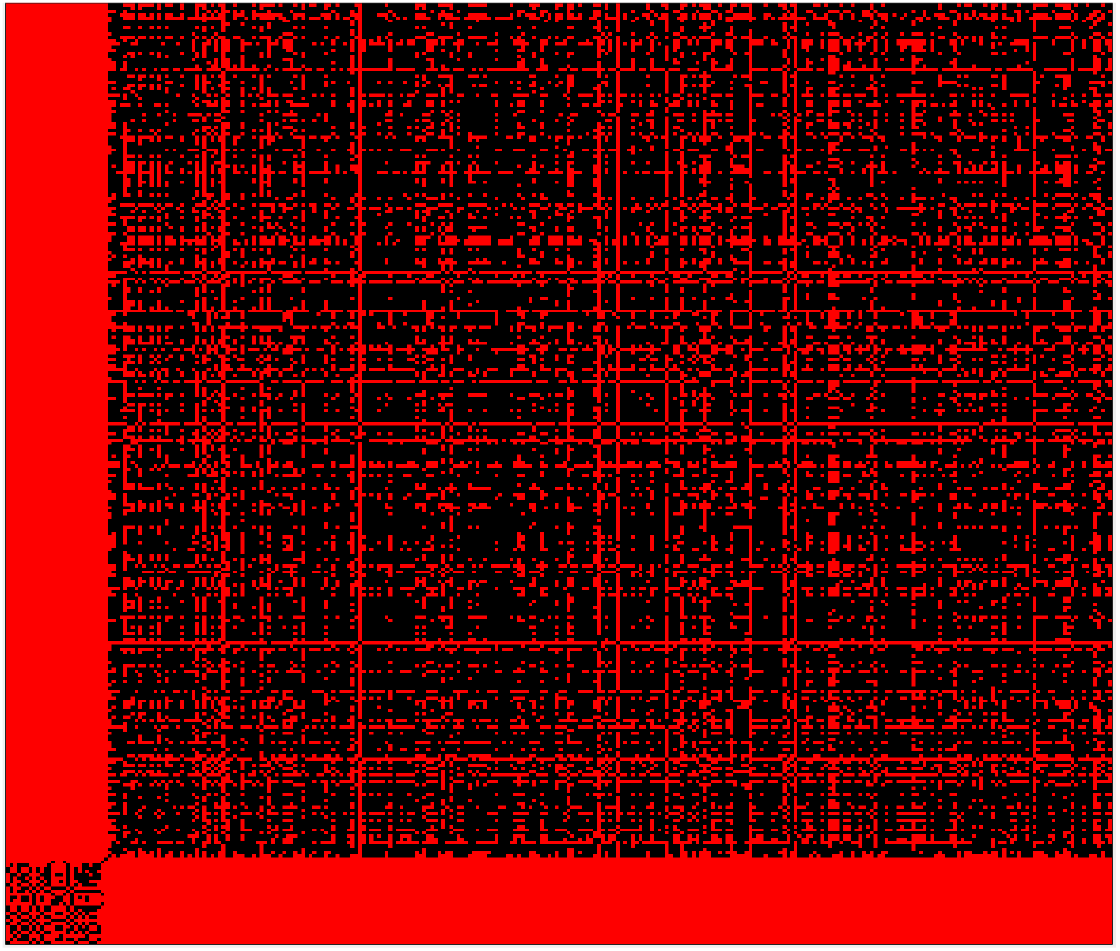
Heat Map Showing Pediatric Cluster with threshold value 0:075

**Figure 3:**
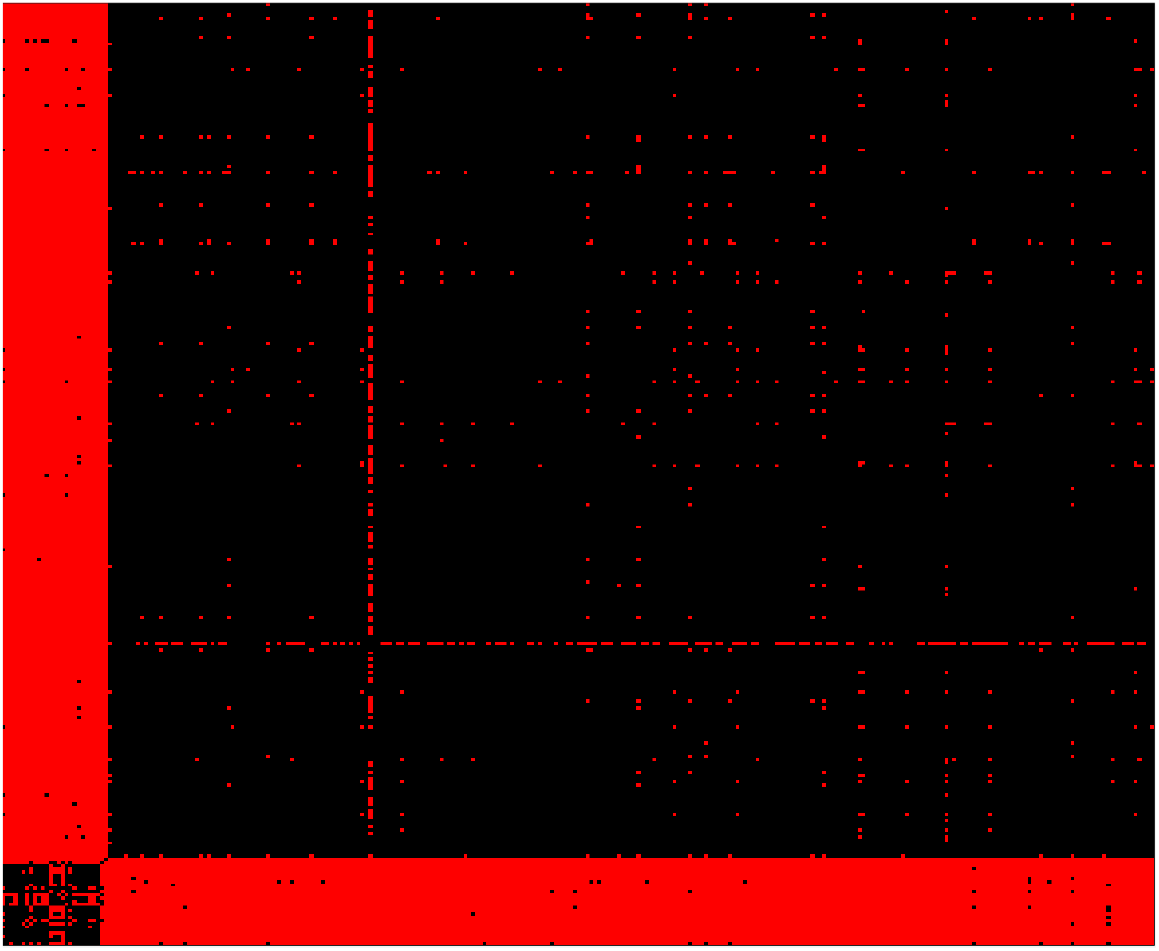
Heat Map Showing Pediatric Cluster with threshold value 0:1

**Figure 4:**
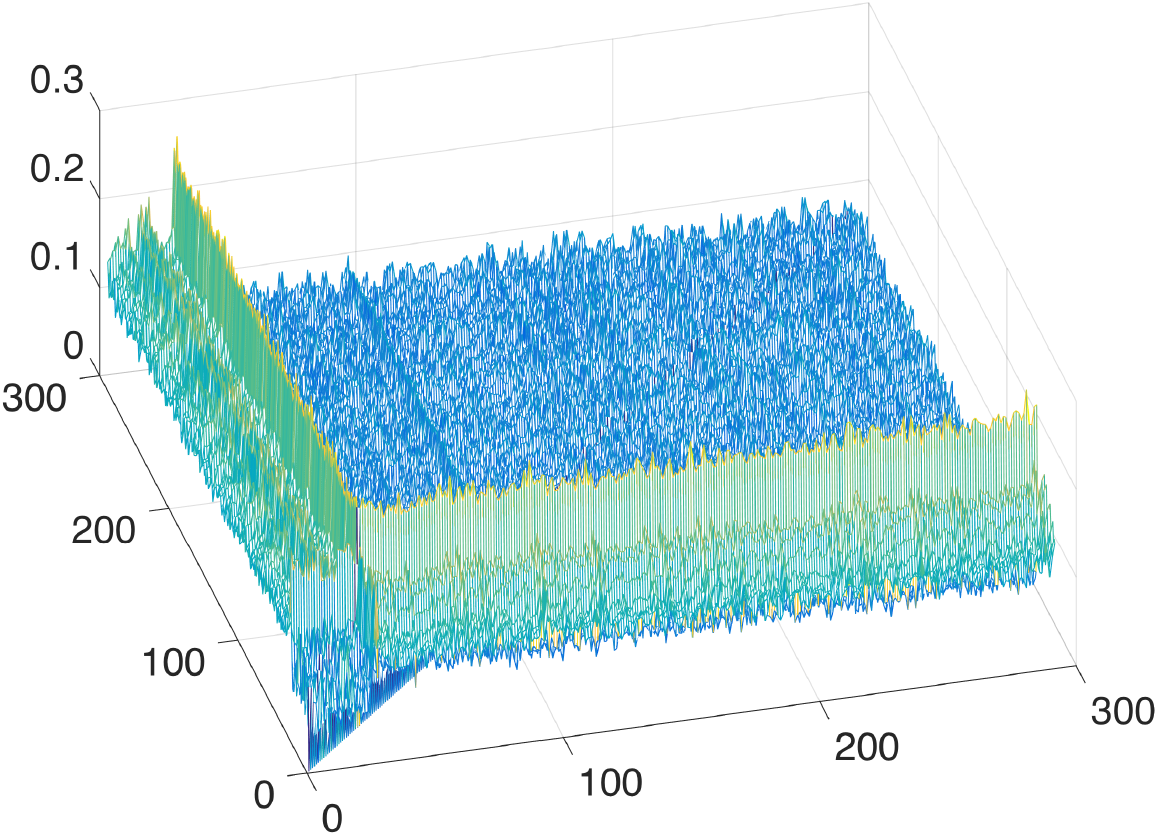
3D Plot Showing Pediatric Cluster

**Figure 5:**
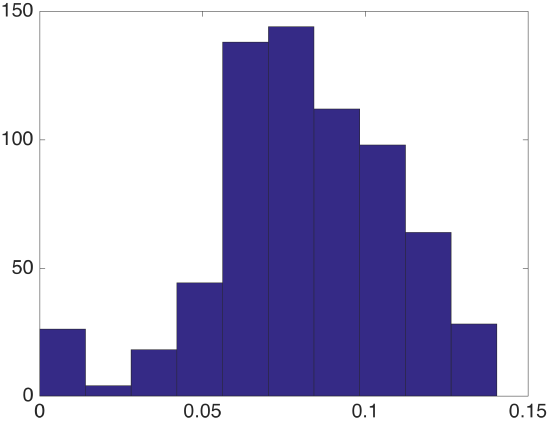
Distances within the Pediatric Cluster

**Figure 6:**
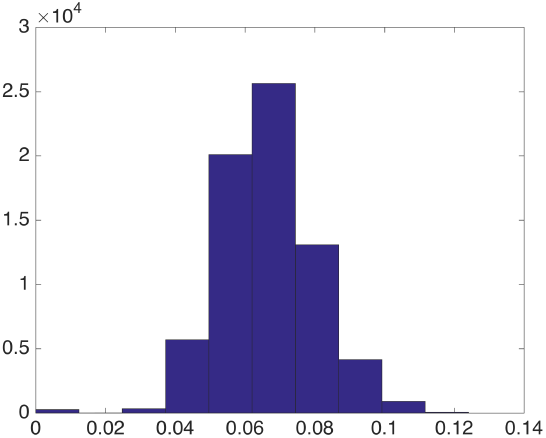
Distances within the Adult Cluster

**Figure 7:**
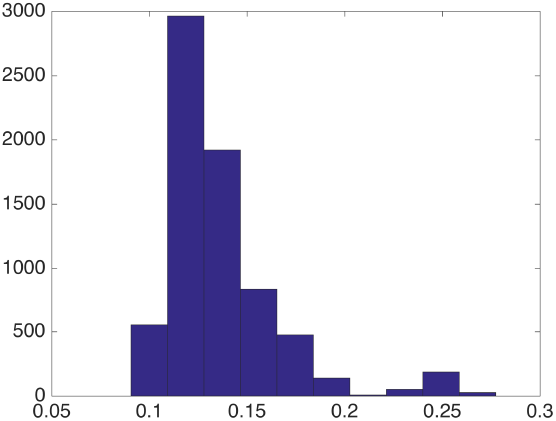
Distances between Pediatric Cluster and Adult Cluster

**Figure 8:**
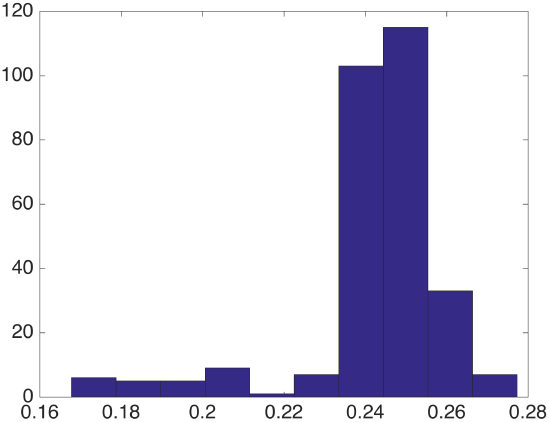
Distances between the Outlier (PIP13-81192) and the other Samples

## 3 Discussion

Sarcomas represent a heterogeneous group of malignant solid tumors of connective tissue. Sarcomas comprise approximately 1.5% of all malignant tumors diagnosed in adults and over 7% of cancers in children [7]. Although the diversity of sarcoma subtypes can be encountered across the age spectrum, there exists a pattern of sarcoma subtypes that significantly distributes between adults and children. For example, osteosarcoma and Ewing sarcoma (malignant bone tumors) are predominant in children and early adults, whereas undifferentiated pleomorphic sarcoma (previously called malignant fibrous histiocytoma), liposarcoma and leiomyosarcoma are extremely rare in children [16, 8].

In addition to the observation that particular sarcoma subtypes predominate in either childhood or in adulthood, there are also differences in the clinical outcomes of adult and childhood sarcoma patients that extend beyond the differences in treatment regimens between adult and childhood sarcomas [1, 2, 8, 21]. With the emergence of next-generation sequencing technologies, we are afforded the opportunity to evaluate the biologic differences between pediatric-associated and adult-associated sarcomas.

In our analysis of 27 sarcoma cases treated at CUMC, only 26 of the 27 original cases would be categorized as a pediatric-associated sarcoma. Interestingly, one case originally included in the pediatric set segregated as an outlier. This case represents a 25 year old female with a history of multiply relapsed, metastatic alveolar soft part sarcoma (ASPS). ASPS is a rare sarcoma subtype comprising 0.2-0.9% of all soft tissue sarcomas [11]. ASPS is extremely rare in childhood, and is more commonly diagnosed in adolescence and young adulthood (15-35 years of age) [7].

A second adult case included in the pediatric cohort is from a 38 year old male with metastatic synovial sarcoma. In contrast to the previous adult cases of ASPS, this case segregated with the pediatric cohort. Synovial sarcoma is a soft tissue sarcoma with a peak incidence in the 3rd decade of life, and with about 1/3 of cases occurring within the pediatric age range [23]. Synovial sarcoma is more common than ASPS and is the most frequent non-rhabdomyosarcomatous soft tissue sarcoma in adolescents and young adults [28]. Although historical differences in the approach to therapy between pediatric and adult oncologists have existed for the treatment of sarcomas and other tumors, there has been acknowledgement in the adult oncology community of the clinical utility of pediatric-based regimens for the treatment of sarcomas occurring in adulthood [5, 22]. However, despite use of more dose-intense chemotherapeutic approaches to the treatment of sarcomas in adulthood, pediatric-associated sarcomas diagnosed and treated in adulthood continue to have inferior outcomes compared to treatment in childhood [9, 10].

These observations suggest that there may exist age-dependent differences in the biology of sarcomas. However, it is unclear what the thresholds for age may be that would contribute to differential responses to treatment and clinical outcome as the cutoffs for age and the definition of “adult age” has varied in the literature. The results from this analysis suggest that the sarcoma subtype may supersede, in this instance, the contribution of age to the biologic behavior and genomic signature. So from this classification scheme, it seems that there are indeed biologic differences between sarcoma subtypes that are generally associated with childhood (such as synovial sarcoma) versus those more commonly associated with adulthood (such as ASPS), and provides a rationale for the use of pediatric regimens for the treatment of these diseases regardless of the patient’s age.

Genomic characterization of a larger cohort of pediatric-associated and adult-associated sarcomas will be imperative in specifically clarifying the genomic lesions that result in the clinical differences in behavior of sarcomas across the age spectrum. In any event, we did manage to cluster 26 out of the 27 CUMC cases from the TCGA data using our methodology.

We should finally note that the pipeline sketched in Figure 1 is quite general and may be quite useful in clustering various biological networks. These typically may be represented as weighted graphs, and thus after suitable normalization as Markov chains for which there exist the corresponding stationary measures. Optimal mass transport theory realized by the Earth Mover’s Distance seems to be an ideal tool for capturing distances among these measures, and thus leads to a natural clustering/classification framework.

## 4 Methods

### 4.1 Overall Sketch

Figure 9 illustrates the overall pipeline of the clustering methodology described in the previous sections. The basic idea is that once one has defined the network topology (in this case via the Human Protein Reference Database), and the weights connecting the nodes (derived here from the mass action principle), one can use in a straightforward manner an invariant of each network, and then compute the distance matrix defined by the EMD or Wasserstein 1-metric. In the next section, we will review the definition and properties of this central mathematical object underpinning our analysis.

**Figure 9:**
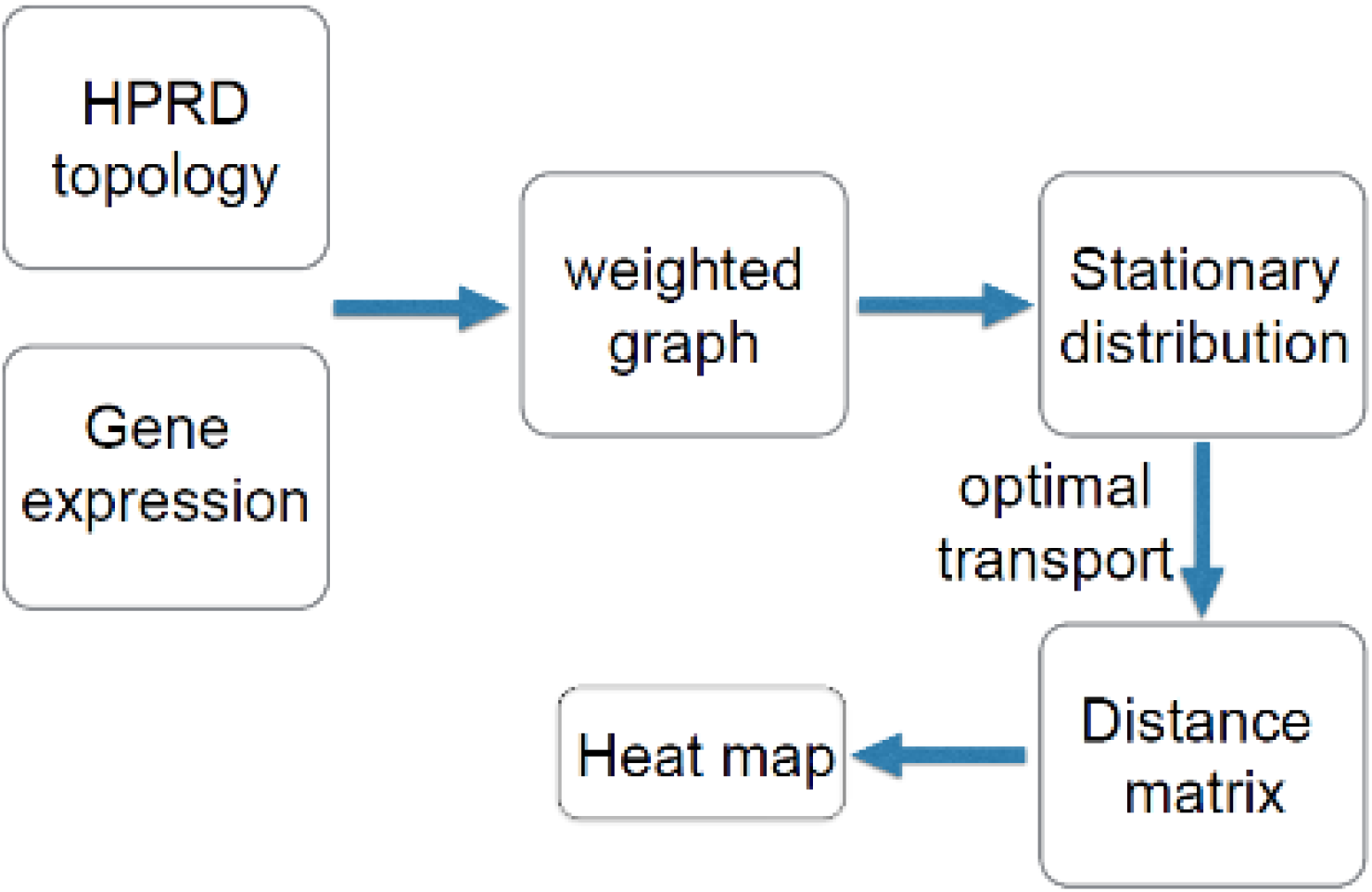
Overall Sketch of Method.

### 4.2 Earth Mover’s Distance

In this section, we briefly review the Earth’s Mover’s Distance (EMD) from optimal mass transport theory, the key method on which all the previous results were based. The classical Earth Mover’s Distance was formulated by Monge in 1781 to solve the problem of moving a pile of soil to a excavation site with the least amount of work relative to some cost. See Figure 10. For full details and a long lists of references, see the monographs [19, 26, 27].

**Figure 10:**
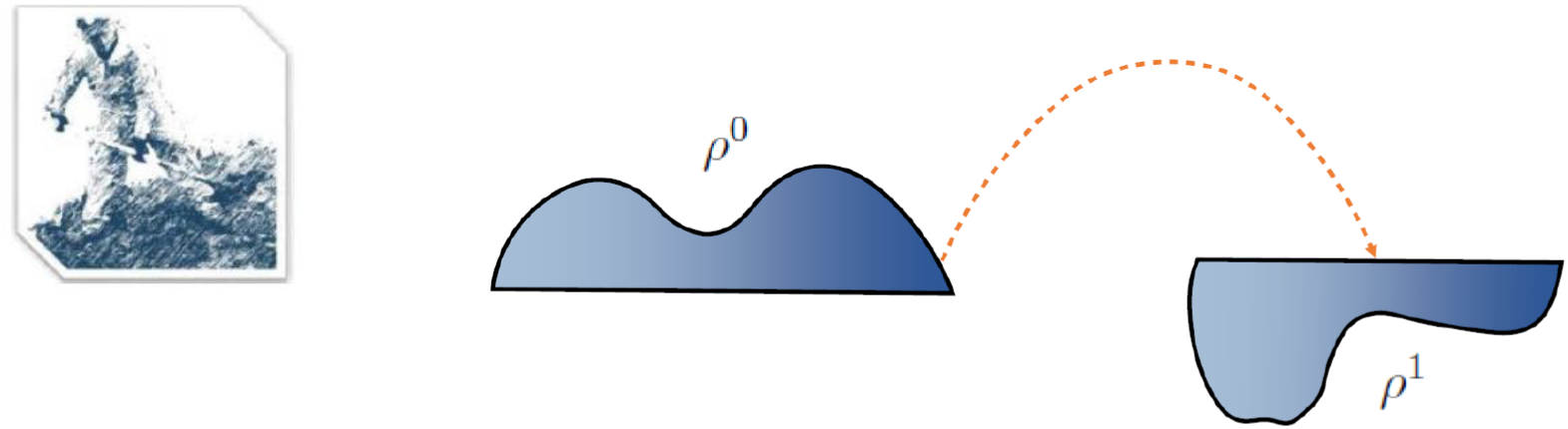
Classical Earth Mover’s Problem.

Mathematically, we let *ρ*^0^ and *ρ*^1^ denote two probability densities on ℝ ^*m*^. This means that *ρ^i^*: ℝ ^*m*^ → ℝ with *ρ^i^* ≥ 0 for *i* = 0,1, such that

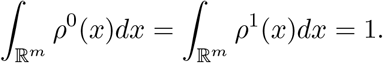

Then the *Earth Movers’ Distance* (also called the *Wasserstein 1-metric, W*_1_) between them is

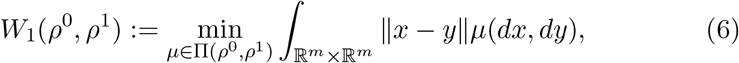

where ∏( *ρ*^0^, *ρ*^1^) denotes the set of couplings between *ρ*^0^ and *ρ*^1^. The Wasserstein-1 distance has the dual formulation [6]

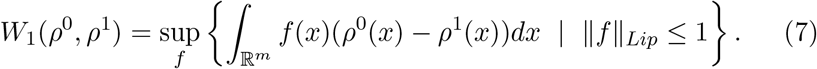

Here

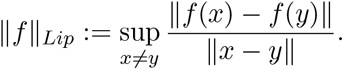

Clearly when *f* is differentiable, ||*f* ||_***Lip***_ ≤ 1 is equivalent to ||∇_***x***_*f* || ≤1. So formally, the above can be rewritten as

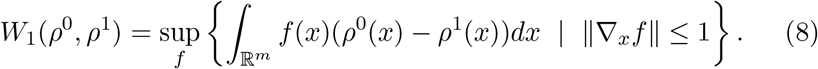

One can then take the dual once again, i.e., starting from (8), one sees that

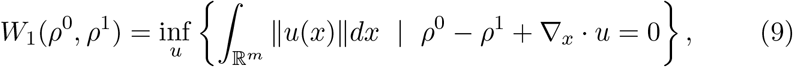

of *W*_1_ with flux u being the optimization variable.

As described above, using this “dual of the dual” formulation applied to sparse graphs, one gets a tremendous saving in computational burden since equation (6) involves solving systems on the order of the square of the number of nodes, while equation (9) is of the order of the number of edges.

## Acknowledgements

This project was supported by AFOSR grants (FA9550-15-1-0045 and FA9550- 17-1-0435), grants from the National Center for Research Resources (P41- RR-013218) and the National Institute of Biomedical Imaging and Bioengineering (P41-EB-015902), National Institutes of Health (1U24CA18092401A1), and a postdoctoral fellowship through Memorial Sloan Kettering Cancer Center.

